# Microclimate buffering and thermal tolerance across elevations in a tropical butterfly

**DOI:** 10.1101/2019.12.19.882357

**Authors:** Gabriela Montejo-Kovacevich, Simon H. Martin, Joana I. Meier, Caroline N. Bacquet, Monica Monllor, Chris D. Jiggins, Nicola J. Nadeau

**Affiliations:** Department of Zoology, University of Cambridge, Cambridge, CB2 3EJ, UK; Institute of Evolutionary Biology, The University of Edinburgh, Edinburgh, EH9 3FL, UK; St John’s College, University of Cambridge, Cambridge, CB2 3EJ, UK; Universidad Regional Amazónica de Ikiam, Tena, Ecuador; Animal and Plant Sciences, University of Sheffield, Sheffield, S10 2TN, UK

## Abstract

Microclimatic variability in tropical forests plays a key role in shaping species distributions and their ability to cope with environmental change, especially for ectotherms. Yet, currently available climatic datasets lack data from the forest interior and our knowledge of thermal tolerance among tropical ectotherms is limited. To tackle this, we studied natural variation in the microclimate experienced by a tropical genus of butterflies (*Heliconius* sp.) along their Andean range across a single year. We found that the forest strongly buffers temperature and humidity in the understory, especially in the lowlands where temperatures are more extreme. There were systematic differences between our yearly records and macroclimate databases (WorldClim2), with lower interpolated minimum temperatures and maximum temperatures higher than expected. We then assessed thermal tolerance of ten *Heliconius* butterfly species in the wild and showed that populations at high elevations had significantly lower heat tolerance than those at lower elevations. However, when we reared populations of the widespread *H. erato* from high and low elevations in a common-garden environment, the difference in heat tolerance across elevations was reduced, indicating plasticity in this trait. Microclimate buffering is not currently captured in publicly available datasets but could be crucial for enabling upland shifting of species sensitive to heat such as highland *Heliconius*. Plasticity in thermal tolerance may alleviate the effects of global warming on some widespread ectotherm species, but more research is needed to understand the long-term consequences of plasticity on populations and species.

**Summary statement:** Tropical forests along the Andes were found to greatly buffer climate. The butterflies inhabiting high elevations were less thermally tolerant but not when reared in common-garden conditions, indicating plasticity.

## Introduction

Land-use and climate change are forcing organisms in the Anthropocene to move, adapt or die (Dirzo et al., 2014). But moving in an increasingly fragmented landscape or adapting to an everchanging climate might be difficult for organisms usually exposed to a narrow range of environmental conditions. Those restricted to stable climates or with limited dispersal abilities have been predicted to be at particular risk of extinction (Bestion et al., 2015; Kingsolver et al., 2013). Despite tropical ectotherms making up half of the animal species described, our knowledge of their potential to cope with high temperatures in natural settings is limited, especially along elevational clines (García-Robledo et al., 2016; Sheldon, 2019). We therefore need a better understanding of the ability of ectotherms to cope with temperatures across elevations and of the climate buffering potential of tropical forests (García-Robledo et al., 2016; Sheldon, 2019).

Since the 1960’s, the notion that “mountain passes are higher in the tropics” (Janzen, 1967) has inspired generations of ecological and evolutionary research. Janzen’s ‘seasonality hypothesis’ predicts that the reduced seasonality in the tropics selects for narrower thermal tolerances than in temperate zones, which would in turn limit their dispersal across elevations (Angilletta, 2009; Nadeau et al., 2017; Sheldon et al., 2018). Subsequent empirical studies have shown that thermal breadth of insects is indeed higher in temperate zones, where seasonality is stronger than in the tropics (Deutsch et al., 2008; Shah et al., 2018). Furthermore, the great level of specialisation of tropical montane species, reflected by high levels of endemism and beta diversity at high altitudes, may highlight further temperature specificity, and therefore susceptibility to the effects of global warming (Polato et al., 2018; Sheldon, 2019). However, in the face of climate and land-use change in lowland habitats, mountains can act as refugia. Some vulnerable lowland organisms are already shifting their ranges upward (Lawler et al., 2013; Morueta-Holme et al., 2015; Scriven et al., 2015), it is critical to ascertain the potential of ectotherms to overcome the physiological barriers that mountains pose.

Janzen’s hypothesis has often been tested at the macroecological scale and with interpolated data from weather stations, assuming that tropical ectotherms live at ambient air temperature (Pincebourde and Suppo, 2016). However, this ignores the microclimate differences most relevant to organisms inhabiting tropical forests (Potter et al., 2013). Tropical forests are very heterogenous habitats with a particularly steep vertical climatic gradient, such that the understory is often more than 2°C cooler than the canopy and spanning an 11% difference in relative humidity (Scheffers et al., 2013). This complexity is not fully captured by interpolated datasets often used in ecological modelling (e.g. WorldClim2, Fick and Hijmans 2017), with mean temperatures in some cases overestimated by 1.5°C (Blonder et al., 2018; Kearney and Porter, 2009; Storlie et al., 2014). Thus, the biological relevance of studies in the tropics using weather station data is limited, as they are positioned specifically to minimise habitat characteristics that can be crucial in determining the thermal tolerance of local organisms (Frenne and Verheyen, 2016; Jucker et al., 2018; Senior et al., 2017). These biases could become even more pronounced at higher elevations, where weather stations are very sparse in the tropics (Fick and Hijmans, 2017; Paz and Guarnizo, 2019).

Extreme climatic events and increased daily climatic ranges may be more important determinants of the biological responses to climate change than temperature mean alone (Sheldon and Dillon, 2016). However, microclimates can buffer ambient temperatures and might act as refugia against such extremes. This buffering could encourage shifts in forest vertical stratification across elevations, such that at higher elevations species may become more arboreal thanks to cooler canopies (Scheffers et al., 2014). However, highly mobile ectotherms such as flying insects often need to reach food sources hundreds of meters apart and across different forest layers, such that behavioural buffering might not be possible. Furthermore, ectotherms can have vastly different ecologies through different life-stages, both in the microclimates of the forest they inhabit and in their ability to cope with thermal extremes (Klockmann et al., 2017; MacLean et al., 2016; Pincebourde and Casas, 2015; Steigenga and Fischer, 2009). Therefore, the fate of ectotherms in tropical forests will depend largely on their own thermal tolerances, as well as on the availability of local climate refugia that buffer against extreme temperatures (Nowakowski et al., 2018; Pincebourde and Casas, 2015; Scheffers et al., 2014).

Plasticity and evolutionary potential in thermal tolerance could help ectotherms cope with human-induced climate and habitat change (Hoffmann and Sgrò, 2011). Tropical species are predicted to have evolved reduced thermal plasticity compared to temperate species, due to low or absent seasonality (Sheldon et al., 2018; Tewksbury et al., 2008). A recent review found that ectotherms, in general, have low thermal tolerance plasticity, with most species having less than a 0.5°C acclimation ability in upper thermal limits (Gunderson and Stillman, 2015; Sheldon, 2019). Detecting evolutionary change in the wild is challenging, especially in the tropics where long-term monitoring schemes are extremely rare (Merilä and Hendry, 2014). However, in two tropical *Drosophila* species, moderate levels of desiccation stress have led to adaptive evolutionary responses in lab conditions (van Heerwaarden and Sgrò, 2014), whereas flies were not able to track the changes with higher and unrealistic levels of desiccation stress (Hoffmann et al., 2005; Sheldon, 2019). Thus, a better knowledge of the plasticity and evolutionary potential of thermal tolerance in tropical insect species will be key to predicting their ability to cope with the warming climate, and tests with realistic levels of environmental change are required.

Accurately predicting the responses of tropical ectotherms to climate and land-use change therefore requires that we understand two complementary aspects of the system: the thermal and humidity buffering potential of tropical forests across altitudinal gradients and the thermal tolerance of the organisms inhabiting them. In this study we (i) measured microclimates for a full year (temperature and humidity) across the elevational range of *Heliconius* butterflies and assess the accuracy of publicly available climatic predictions for the same locations, (ii) tested heat tolerance of ten butterfly species in the wild and (iii) reared offspring from high and low altitude populations of *H. erato* in common-garden conditions to test whether differences observed between wild populations were genetic or plastic.

## Methods

### STUDY SYSTEM AND WILD BUTTERFLY COLLECTION

We collected high (mean=1398 meters above sea level, m.a.s.l.) and low (mean=495 m.a.s.l.) altitude populations of *Heliconius* butterflies with hand nets along the western and eastern slope of the Ecuadorian Andes, at a similar latitude (Supplementary Information, Fig. S1). Every *Heliconius* species encountered at each site was collected, but only those species with more than 5 individuals at each elevation (high and/or low) were included in the analyses (15 out of 329 wild individuals removed). Detached wings were photographed dorsally and ventrally with a DSLR camera with a 100 mm macro lens in standardised conditions, and wing area was measured with an automated pipeline in the public software Fiji (following Montejo-Kovacevich et al., 2019). All the images are available in the public repository Zenodo (https://zenodo.org/communities/butterfly/) and full records with data are stored in the EarthCape database (https://heliconius.ecdb.io). A list with all the individuals and links to both databases can be found in the Appendix 1.

### MICROCLIMATES ACROSS ALTITUDES

Microclimates across elevations on both sides of the Ecuadorian Andes were characterised by recording temperature and relative humidity every hour in the understory and mid-layers of the forest for a full year (Fig. S1), between February 2017 and February 2018. We used 40 HOBO temperature loggers (model: HOBO UA-001-08 8K; accuracy: 0.5°C, resolution: 0.1°C, 10 per area) and 16 high accuracy humidity and temperature HOBO data loggers (model: HOBO U23-001; temperature accuracy: 0.21°C; temperature resolution: 0.02°C; relative humidity accuracy: 2.5%; relative humidity resolution: 0.05%, four loggers per area). We chose 28 accessible forest sites that had not recently been disturbed by humans, usually inside or near nature reserves (localities in Table S1). Seven of these were at high altitude (mean= 1214 m.a.s.l.) and seven at low altitude (mean= 444.6 m.a.s.l.) in the eastern and western slopes of the Andes (Fig. S1). Sites were at least 250m apart from each other and in the same areas where *Heliconius* populations were sampled for this study. We placed one logger in the understory (mean height= 1.16 m) and one in the subcanopy (mean height= 10.7 m) at each site. Subcanopy loggers were as close as possible to directly above the understorey loggers, and both were hung from tree branches with fishing line and suspended mid-air (Supplementary Video 1). To prevent exposure to direct solar radiation, temperature data loggers were secured inside a white plastic bowl and humidity data loggers between two flat white plastic plates, allowing for horizontal air flow.

### HEAT TOLERANCE

Heat stress resistance of wild-caught individuals was measured with a heat knockdown assay (Huey et al., 1992; Sørensen et al., 2001). Butterflies were tested less than 12h after they were collected in the field and at approximately the same altitude. Individuals were stored in envelopes with damp cotton and fed a small amount of sugary water to standardise hydration levels before being tested. Butterflies were placed in individual glass chambers, fitted with an instant read digital thermometer (accuracy: 1.0°C, resolution: 0.1°C). Glass chambers contained 150g of metal beads to add weight, covered with a Styrofoam platform for butterflies to stand on. Five chambers were introduced at a time into a plastic hot water bath at 51°C, and we recorded the time until the interior of the chamber, where the butterflies are placed, reached 39.0°C (“heating up time” hereafter). For the duration of the assay the temperature inside the chambers was kept between 39.0°C and 41.0°C, by increasing or decreasing the hot water bath temperature as required. Tests in which the chamber temperature went below this range during the duration of the assay were removed from further analyses. We recorded the time and exact temperature at heat knock-down, defined by the loss of locomotor performance of the individual butterfly (Huey et al., 1992), i.e. when the butterfly’s legs collapsed or it fell on its side. Temperature at KO was accounted for in the models, as in 89 out of 496 assays the chamber went above 41°C (Fig. S6), but assays in which temperature went above 41.9°C were removed from all analyses (n=2). Butterflies were monitored for a maximum of 60 minutes, and this maximum value was used for those that had not been knocked out within the timeframe (n=1).

### COMMON-GARDEN REARING

Fertilised females of *H. erato* were caught in the wild with hand nets in high (mean=1348 m.a.s.l.) and low altitude localities (mean=543 m.a.s.l.), in the vicinity of the data loggers used for microclimates analyses. Females from both altitudes were kept in separate cages of purpose-built insectaries at the Universidad Regional Amazónica Ikiam (Tena, Ecuador, 600 m.a.s.l.). Eggs were collected three times per day and individuals were reared in separate containers throughout development, but stored in the same area of the insectary and randomly assorted. All offspring were individually fed the same host plant, *Passiflora punctata*. Development rates, pupal and adult mass were recorded for all offspring. Common-garden reared offspring were blindly tested for heat knock-down resistance one day after emergence following the same protocol as for wild-caught individuals. Offspring from high and low altitude mothers had individual IDs with no indication of their origin, thus we were able to blindly test five individuals at a time and avoid potential observer bias. Adult offspring wings were photographed and their wing areas measured with an automated pipeline in the open-access software Fiji (following Montejo-Kovacevich et al., 2019).

### STATISTICAL ANALYSES

All analyses were run in R V2.13 (R Development Core Team 2011) and graphics were generated with the package *ggplot2* (Ginestet, 2011). Packages are specified below and all R scripts can be found in the public repository Zenodo (Zenodo: TBC).

#### Microclimates across altitudes

Our data showed low seasonality (standard deviation of monthly averages (Bio4)= 0.63, Note S1), as expected for latitudes near the Equator, therefore it was not subdivided into months. To determine the range of temperatures and humidities that butterfly populations are exposed to at different altitudes and sides of the Andes, we first estimated daily maxima, mean, and minima temperature and humidity per data logger across the year. We used linear mixed effect models (LMMs) to determine temperature differences across forest strata (understory and subcanopy), implemented with lmer() function from the lme4 package (Bates et al., 2015). We included forest strata, altitude and date as fixed effects, and as random effects we placed data logger ID nested within site and area (altitude+side of the Andes). Dates were standardized to a mean of zero and unit variance to improve model convergence (Zuur et al., 2009). Significant differences between forest strata and altitudes were assessed via post-hoc comparisons with Tukey tests). Mean diurnal range was estimated per day per data logger (as the daily minimum subtracted from the daily maximum) and then averaged across the year. To obtain the rate of increase in understory temperature for every 1°C increase in subcanopy temperature we fitted linear models with the hourly data to obtain the relationship between understory temperature and subcanopy temperature (following González del Pliego et al. 2016).

To understand the atmospheric water imbalance of each microclimate and elevation, we calculated vapour pressure deficit (VPD, in hectoPascals - hPa) based on the hourly high-resolution relative humidity and temperature measurements taken across the year. VPD is the difference between how much moisture the air can hold before saturating and the amount of moisture actually present in the air, i.e. a measure of the drying power of the air. VPD relies on both temperature and relative humidity, making it more biologically relevant than relative humidity alone (Bujan et al., 2016). Relative humidity, which is directly measured by our humidity dataloggers, depends partially on air pressure, thus we do not need to further account for it. Vapour pressure deficit is linked to water transport and transpiration in plants, and negatively correlated with survival and growth in trees and with desiccation resistance in ectotherms (Bujan et al., 2016). It is calculated as the difference between saturation water vapour pressure (e_s_) and water vapour pressure (e) (Jucker et al., 2018). Given that relative humidity (RH), which is directly measured by our data loggers, can be expressed as RH=(e/e_s_) and VPD is calculated:

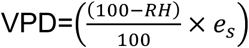

And e_s_ is derived from temperature (in °C) using Bolton’s equation (Bolton, 1980; Jucker et al., 2018)

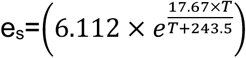

VPD was estimated for every coupled hourly temperature and relative humidity record, i.e. from the same data logger at the same time/date, and we then calculated annual mean VPD (VPD_mean_) and mean daily maximum VPD (VPD_max_) per logger.

#### Comparing measured microclimate with coarse resolution climatic data

To assess the differences between field obtained microclimate data and publicly available ambient climate data we compared our microclimate data to coarser-resolution data from the WorldClim2 database, which is widely used in ecological modelling (Fick and Hijmans, 2017). Worldclim2 climate grid data is generated by interpolating observations from weather stations from across the globe since the 1970s, which are typically located in open environments (Jucker et al., 2018). We extracted climatic data for the mean coordinates from the seven sites of our four study areas (East/West, highlands/lowlands, Fig. S1) with the maximum resolution available (1km^2^, 30 seconds), using the package “raster” (Hijmans et al., 2019). Following a recent study (Jucker et al., 2018), we focused on comparing WorldClim2 interpolated monthly means of daily temperatures, annual mean temperatures (Bio 1) and annual mean diurnal range (Bio 2) to the equivalent bioclimatic predictors calculated with our microclimate data. Thus, we extracted the following WorldClim2 bioclimatic variables for all the areas under study:

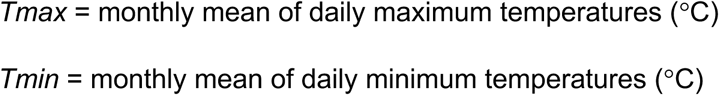

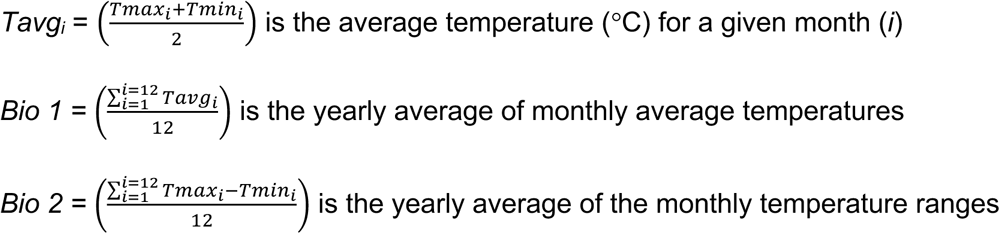

Bio1 is more commonly known as the annual mean temperature and Bio2 as the annual mean diurnal temperature range. The latter is mathematically equivalent to calculating the temperature range for each day in a month and averaging across that month. Since our microclimate data showed low yearly seasonality (temperature SD=0.63)and to avoid biases in months where not all records were available (e.g. when changing logger batteries halfway through the year), we averaged across the whole year.

#### Thermal tolerance across species in the wild and across H. erato common-garden reared families

To test variation in thermal tolerance across species in the wild and across families in common-garden reared conditions, we first used an ANOVA approach, with species or brood as a factor explaining the variation in heat knockdown time. We then estimated within-species and within-brood trait repeatability, or intra-class correlation coefficient (ICC), with a linear mixed model approach. This requires the grouping factor to be specified as a random effect, in this case species (for wild individuals) or brood (for common garden-reared offspring), with a Gaussian distribution and 1000 parametric bootstraps to quantify uncertainty, implemented with the function rptGaussian() in *rptR* package (Stoffel et al., 2017). By specifying species/brood ID as a random effect, the latter approach estimates the proportion of total thermal tolerance variance accounted for by differences between species or families.

To determine the effects of altitude on thermal tolerance in wild butterflies and common-garden reared offspring we fitted two separate linear mixed models (LMMs), implemented with lmer() function from the lme4 package (Bates et al., 2015). Both models initially included heat knock down time as the response variable, and altitude at which the wild individual or mother of the brood were collected, sex, wing size (plus all two-way interactions between them), minutes that the chamber took to reach 39°C, and temperature at knock-out as fixed effects. Additionally, the common-garden reared offspring model included development time (days from larva hatching to pupation) and brood egg number (to control for time the brood mother spent in the insectary) as fixed effects. The random effects for the wild individuals model and for the common-garden reared offspring model, were species identity and brood mother identity respectively. All fixed effects were standardized to a mean of zero and unit variance to improve model convergence (Zuur et al., 2009).

We implemented automatic model selection with the step() function of the lmerTest package (Kuznetsova et al., 2017), which performs backwards selection with likelihood ratio tests and a significance level of α = 0.05, first on the random effects and then on the fixed effects to determine the simplest best fitting model. Model residuals were checked for homoscedasticity and normality. To obtain *P*-values for the fixed effects we used the anova() function from the lmerTest package, which uses Satterthwaite approximation (Kuznetsova et al., 2017). We estimated with the coefficient of determination (R^2^) the proportion of variance explained by the fixed factors alone (marginal R^2^, R^2^_LMM(m)_) and by both, the fixed effects and the random factors (conditional R^2^, R^2^_LMM(c)_), implemented with the *MuMIn* library (Bartón, 2018; Nakagawa et al., 2017).

## Results

### MICROCLIMATE VARIABILITY ACROSS ALTITUDES

Overall, the patterns of differentiation between forest layers and altitudes are very similar across sides of the Andes in our study (Fig. 1 A vs B), and there was low seasonality across months (Fig. 1, Note S1). The lowland sites were, on average, 4.1°C warmer than the highland sites which were over 750m apart in elevation (Table S1, S2). Annual mean temperatures interpolated from WorldClim2 were always closer to subcanopy strata annual means (Fig. S3 A). In contrast, the understory annual mean temperatures were on average 0.5°C lower than WorldClim2 annual mean temperatures (dotted lines Fig. 1 A1-2, B1-2, Table S2). The minimum temperatures were consistently higher in our microclimate data compared to the interpolated monthly minima estimated in WorldClim2, especially for the highlands (lower dotted line, Fig. 1 A2, B2).

**Figure 1.**
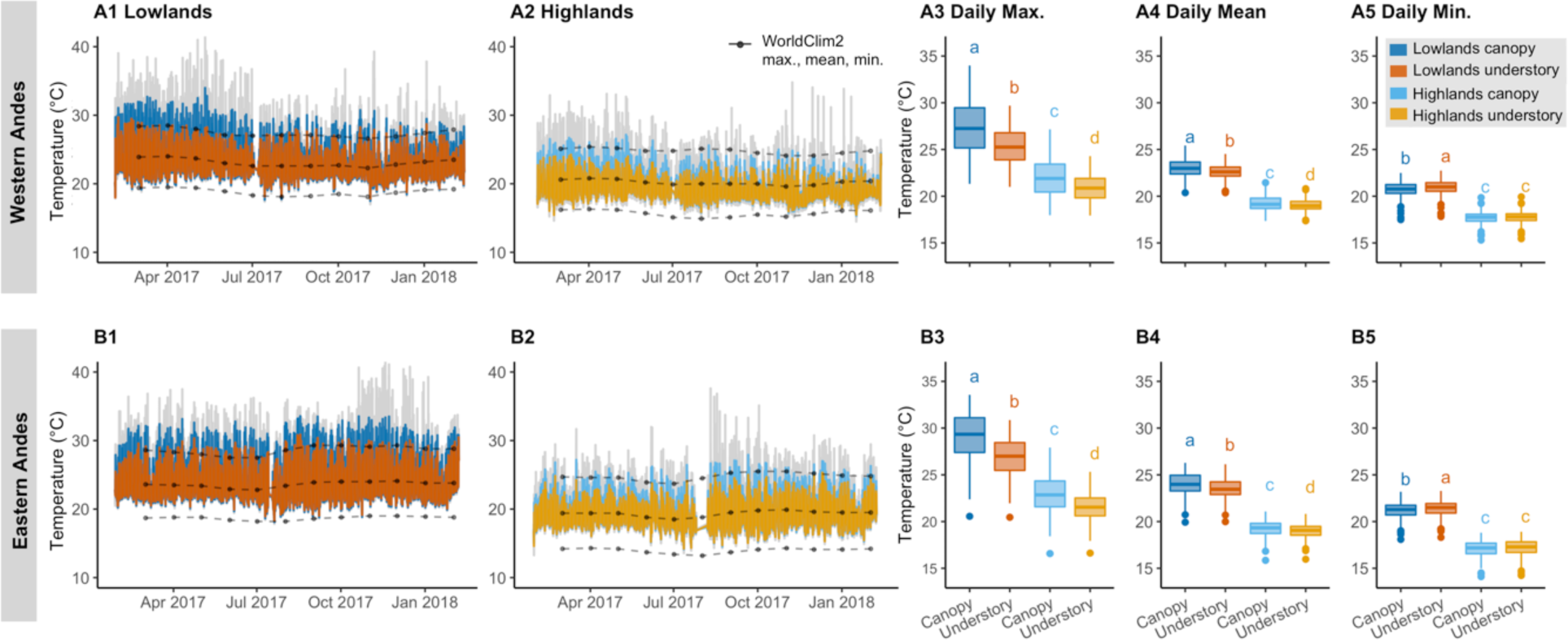
Annual and daily microclimates across forest heights and elevations. Annual microclimate variation recorded every hour (Lowlands/Highland, West/East, A1-2, B1-2), mean daily maximum temperature (A3, B3), mean daily average temperature (A4, B4) and mean daily minimum temperature (A5, B6) from February 2017 until February 2018 in the Western (A1-A5) and Eastern (B1-B5) slopes of the Andes. For A1-2, B1-2 grey lines represent raw data, coloured lines represent hourly temperatures averaged across loggers in each of the four areas and forest layers. For A3-5 and B3-B5 we first obtained individual datalogger daily maximum, mean, minimum temperature, and averaged these to obtain the daily mean values per area/forest layer here plotted. Colours represent microclimates (blues: subcanopy, oranges: understory). Points and dashed lines represent WorldClim2 interpolated monthly maximum (Tmax), mean (Tavg), minimum (Tmin) temperatures for these areas (A1-2, B1-2). The bottom and top of the boxes represent the first and third quartiles, respectively, the bold line represents the median, the points represent outliers, and the vertical line delimits maximum and minimum non-outlier observations. Same superscripts represent no significant differences (Tukey post-hoc test p>0.05).

Forest canopies thermally buffered the understory, but more so during the day and in the lowlands. During the day, the understory thermally buffered the canopy temperature maxima by 1.19-1.62°C in the highlands, and by 1.98-2.11°C in the lowlands (Table 1). At night, understories buffered the temperature minima of the canopies by 0.07-0.12°C in the highlands and by 0.25-0.23°C in the lowlands (Table 1). Thus, the forest buffered high and low temperatures in the understory throughout the day and night, respectively (Fig. 2A). Temperature differences between day and night are greater in the lowlands, where days are warmer (Fig. 2 A), but less so in the understory of all areas. On average, the difference between subcanopy and understory diurnal thermal range in the highlands was 1.34°C, whereas in the lowlands this difference was 2.09°C (Fig. 2 B). However, WorldClim2 interpolations for diurnal thermal ranges were 3.5°C higher than our records in the highlands, resulting in the highlands being predicted to have higher thermal ranges than the lowlands (stars, Fig. 2B). This was the opposite elevational trend to that observed in our data, where thermal ranges were lower in the highlands.

**Table 1.**
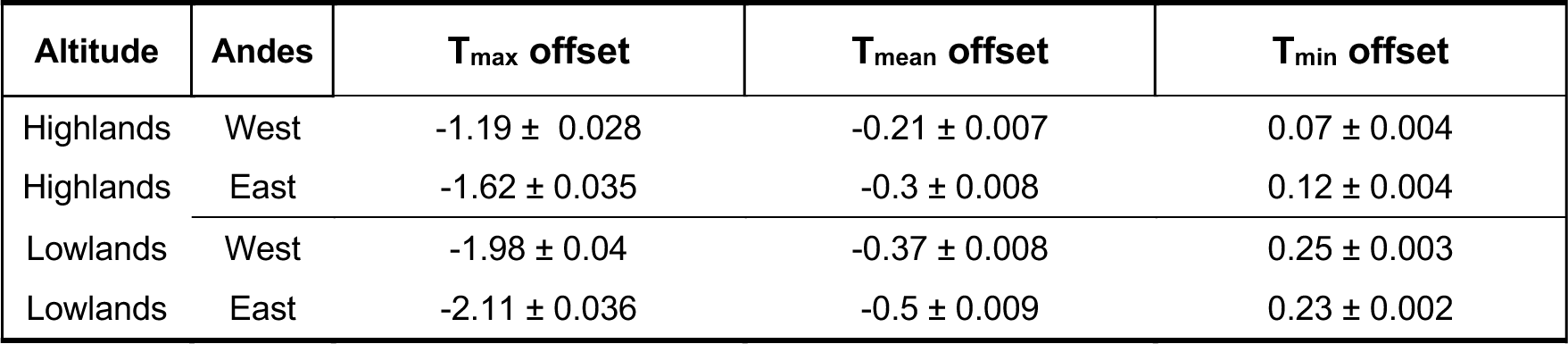
Understory temperature offsets across areas. Offsets were calculated by subtracting subcanopy daily temperatures from understory daily temperatures, thus negative values indicate cooler understories and positive warmer understories. We estimated daily maximum, mean, and minimum temperatures (°C) per datalogger per day, then calculated daily offsets at a given site (pair of loggers) and averaged across the year and dataloggers per area. Values shown are mean and (±) standard error.

**Figure 2.**
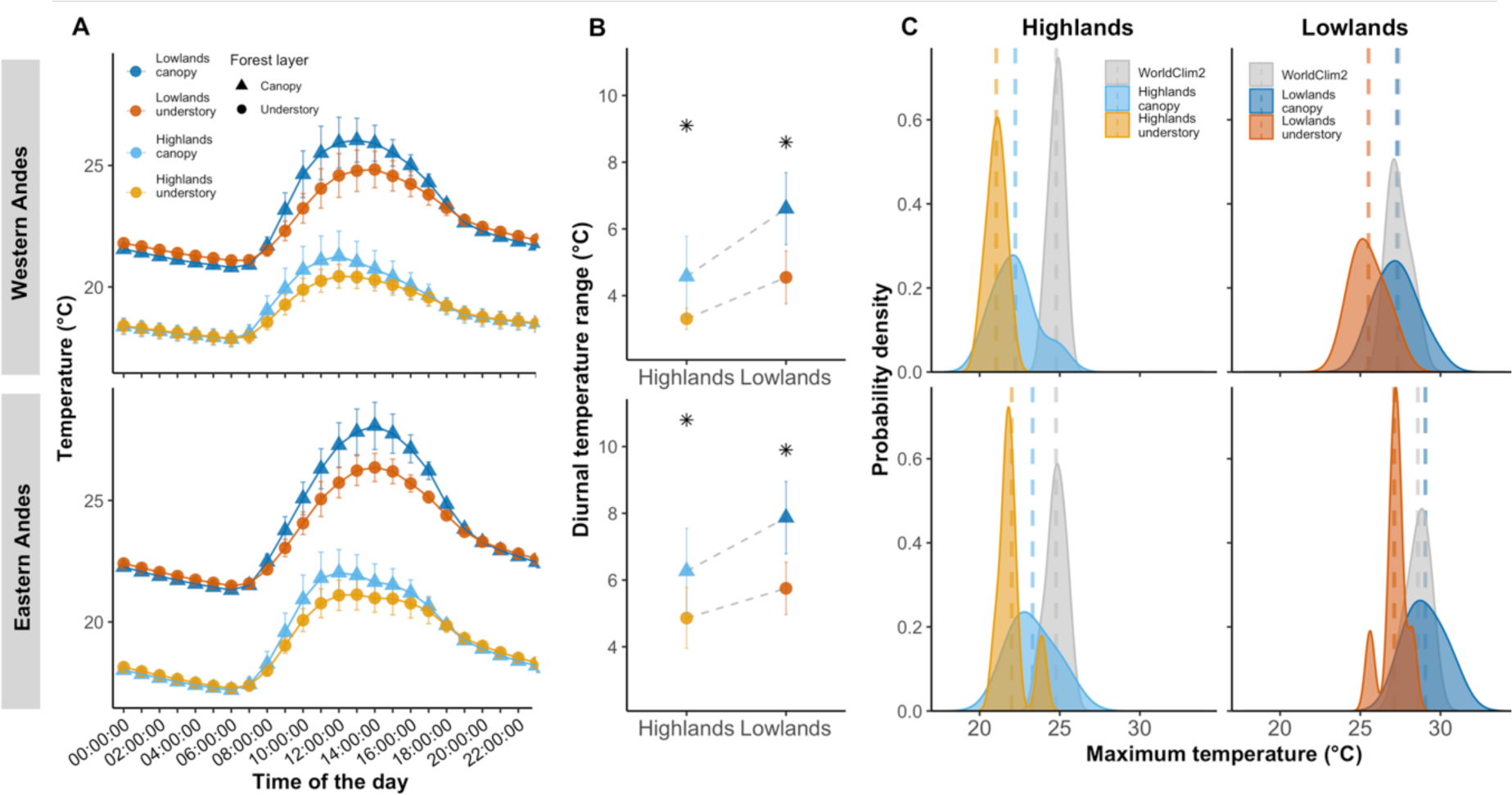
Daily temperature microclimate and interpolated variation. A) Daily temperature variation across altitudes in the Western (top plot) and Eastern (bottom plot) slopes of the Andes, values plotted represent the mean annual temperature across loggers at a given time of the day in one of the four areas (highlands/lowlands, subcanopy/understory). B) Annual diurnal temperature range, calculated for each datalogger across areas and the stars represent the WorldClim2 interpolations for this bioclimatic variable (Bio 2). Error bars represent standard deviation from the mean. C) Mean daily maximum temperature across individual dataloggers for a full year, compared to WorldClim2 interpolated daily maxima (T_max_) for these areas. Vertical dashed lines represent means per group.

The temperature buffering of the lowlands was higher than the highlands, so that for every 1°C increase in subcanopy temperature the understory increased 0.68°C in the lowlands, in contrast to 0.73°C in the highlands (Fig. S2). The lowland canopies exceeded 39°C on 31 days throughout the year, whereas the highlands never did (Fig. 1 A, B). The monthly maximum temperature interpolated by WorldClim2 was close to our measured subcanopy temperature maxima in the lowlands, but 2.01°C higher, on average, in the highlands (Fig. 2C). Wordclim2 monthly minima were also overestimated at both elevations, predicting it to be 2.6°C cooler at night in the highlands and 2.3°C cooler in the lowlands than our measured understory microclimates (Fig. S3 B).

### VAPOUR PRESSURE DEFICIT

In the highlands, the understory daily relative humidity minimum was on average 3.7 percentage points higher than in the subcanopy, whereas in the lowlands the difference was 11.8 percentage points (Fig. S4). The drying power of the air, or vapour pressure deficit (VPD), varied drastically between layers of the forest, but more so in the lowlands (Fig. 3). In the highlands, maximum daily VPD from the same site averaged 1.1 hPa higher in the subcanopy compared to the understory, whereas in the lowlands the difference was 4.8 hPa. The threshold for tree transpiration is thought to be at 12 hPa in the tropical montane areas, above which transpiration, and thus growth, is impeded by the drying power of the air (Motzer et al., 2005). This threshold was exceeded 883 times in the lowland subcanopy across our data loggers, 102 in the lowland understory, 12 times in the highland subcanopy and never in the highland understory (Fig. S5).

**Figure 3.**
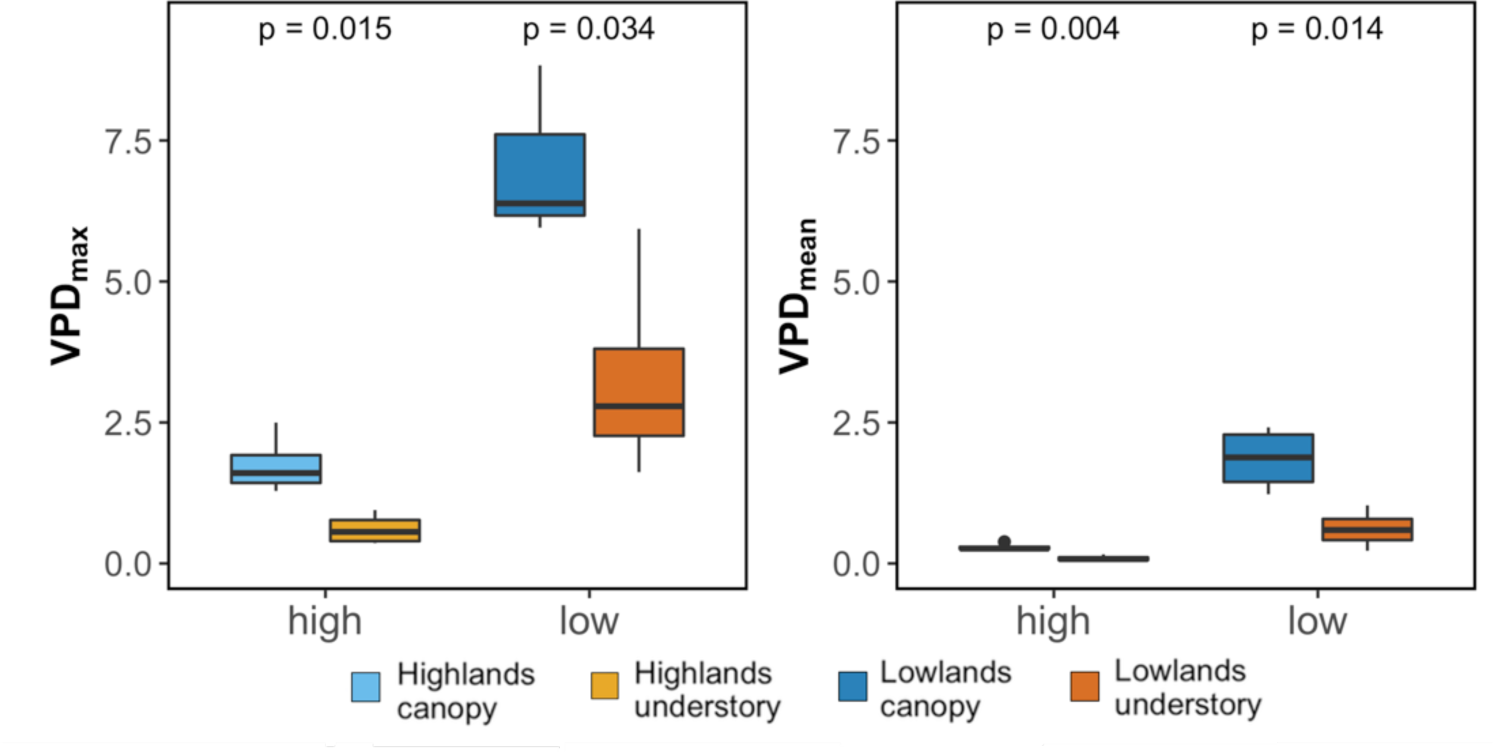
Vapour pressure deficit (VPD, “drying power”, hPa) across microclimates and elevations. A) VPD_max_, mean daily maximum VPD for each datalogger across a year. B) Annual mean VPD (VPD_mean_). P-values are shown for t-tests between subcanopy and understory values.

### HEAT TOLERANCE IN THE WILD

Heat tolerance varied across species (Fig. 4 A, ANOVA: F_9, 268_ = 8.75, P < 0.0001), with 44% of this variation explained by species identity (Repeatability=0.44, S.E.=0.13, P<0.0001). In the highlands, 33% of the individuals tested were knocked out before the chambers reached a temperature of 39°C, in contrast to 7% of the individuals tested in the lowlands (Fig. 4 B, n_high_=60/183, n_low_=6/95). Mean (± standard error) heat tolerance across species was on average 5.4 minutes (±0.57) for highland individuals and 15.9 minutes (±1.43) for lowland individuals (red dashed line Fig. 4 A). Altitude and time until the chamber reached 39°C were significant predictors of knock-out time in wild individuals (Table S1), with the fixed effects alone explaining 29% of the variation in thermal tolerance (R^2^_LMM(m)_=0.29) and 39% when considered together with species identity as a random effect (R^2^_LMM(c)_=0.39). The high-altitude populations of the two most evenly sampled species across altitudes, *H. erato* and *H. timareta*, were less thermally tolerant than their lowland conspecifics (Fig. S7, T-test, *H. erato*: t_77_=-5.3, P<0.0001, *H. timareta*: t_14_=-2.3, p<0.05). In this part of the eastern Andes of Ecuador, the lowland *H. melpomene* have been largely replaced by a cryptic subspecies of *H. timareta* (Nadeau et al., 2014), which explains the low numbers of lowland individuals of *H. melpomene*, a species with a very wide range across the Neotropics.

**Table 2.** Thermal tolerance model summaries. Wild individuals of ten species and common-garden reared *H. erato* final models. Non-significant two-way interactions are not included in this table. Chi.sq, the value of the chi-square statistics. Chi Df, the degrees of freedom for the test. *P*, the *P*-values of fixed and random effects. Sum.Sq, sum of squares. Df, degrees of freedom based on Sattherwaithe’s approximations. *F*, *F*-value. KO, knockout. Egg number per broods, number of eggs previously laid by that brood mother. Times are in minutes.

| Model | N | Response | Random effects |  |  |  | Fixed |  |  |  |  |
| --- | --- | --- | --- | --- | --- | --- | --- | --- | --- | --- | --- |
|  |  |  | Variable | Chi.sq | Chi Df | *P* | Variable | Estimate | df | t-value | *P* |
| Wild individuals | 213 | KO time | Species ID | 8.53 | 1 | 0.003 | Altitude | -4.45 | 151.9 | -5.50 | <0.001 |
|  |  |  |  |  |  |  | Heating up time | 5.76 | 205.2 | 7.87 | <0.001 |
| Common-garden reared | 260 | KO time | Mother ID | 9.43 | 1 | 0.002 | Heating up time | 21.67 | 184.2 | 8.85 | <0.001 |
|  |  |  |  |  |  |  | Egg number (per brood) | 3.04 | 173.7 | 9.80 | <0.001 |
|  |  |  |  |  |  |  | Temperature at KO | 3.12 | 162.9 | 6.02 | <0.001 |
|  |  |  |  |  |  |  | Development (days) | 0.99 | 228.9 | 3.76 | 0.02 |
|  |  |  |  |  |  |  | Altitude*egg number | -0.40 | 224.8 | -2.25 | 0.03 |
|  |  |  |  |  |  |  | Altitude | -1.03 | 134.2 | -2.24 | 0.07 |

**Figure 4.**
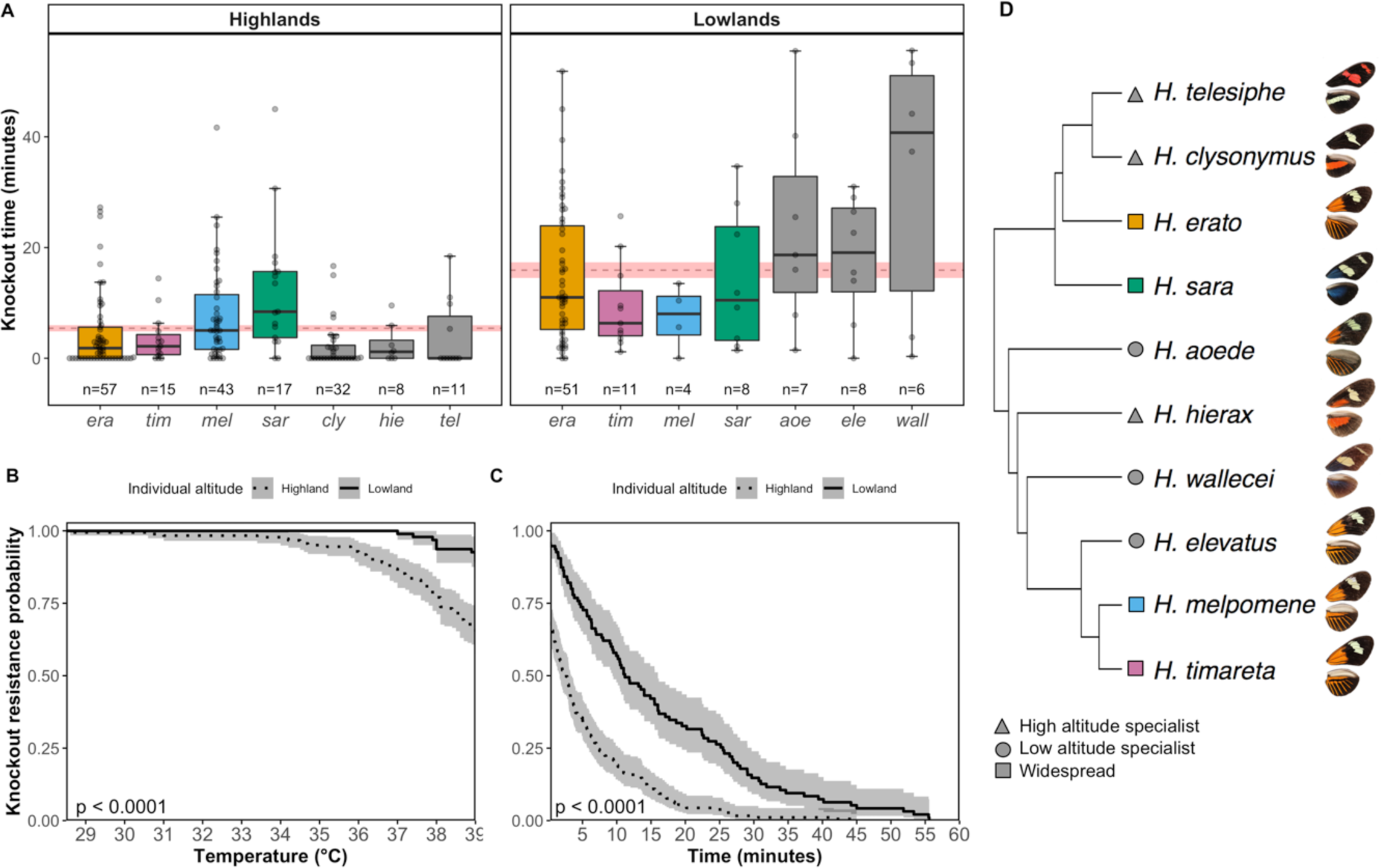
Wild *Heliconius* thermal tolerance. A) Heat knockdown time in minutes across wild individuals from ten *Heliconius* species, coloured species have wide altitudinal ranges whereas grey are high/low altitude specialists. The red dashed line represents the mean heat tolerance of all individuals (regardless of species) at each altitude and the shaded area represents the standard error of the mean. B) Proportion of wild individuals from high and low populations (dotted and solid lines, respectively) that resisted knockout before reaching the experimental temperature of 39°C and C) throughout the heat knockdown experiment (temperatures 39-41°C). Error bars and shaded areas represent 95% confidence intervals of the means and sample sizes for each species are indicated above their label. P is p-value for log-rank test comparing the curves. D) Study species phylogeny (Kozak et al., 2015) and representative images of wings (not to scale).

### HEAT TOLERANCE IN COMMON-GARDEN REARED OFFSPRING

Common-garden reared offspring of *H. erato lativitta* varied in heat tolerance across families (ANOVA, F_14, 262_ = 5.15, P < 0.0001), and 25% of this variation was explained by brood identity (Repeatability=0.25, S.E.=0.10, P<0.0001). In the wild, low-altitude *H. erato lativitta* were, on average, able to withstand high temperatures for ten more minutes compared to high-altitude populations (Fig.4 left, T-test: t_77_=-5.3, P<0.0001). In contrast, when reared in common-garden conditions, individuals from lowland broods were able to withstand heat for only 1.4 minutes longer than offspring from highland broods (Fig.4 right). As a consequence, parental altitude only had a marginally significant effect on offspring thermal tolerance (knock-down time), whereas experimental variables, such as time until the chamber reached 39°C and the temperature at knock-out, were significant predictors of knock-out time in the offspring (Table 1, “Common-garden reared model”). Fixed effects alone explained 39% of the variation in thermal tolerance (R^2^_LMM(m)_=0.39) and 48% when together with brood identity as the random effect (R^2^_LMM(c)_=0.48), indicating trait heritability. The variance in thermal tolerance in wild populations was higher than in common-garden reared offspring (Fig. 5). The number of eggs a mother had previously laid had a positive and significant effect on adult thermal tolerance, and interacted with parental altitude, likely due to high altitude mothers living longer in the insectary (Fig. S8).

**Figure 5.**
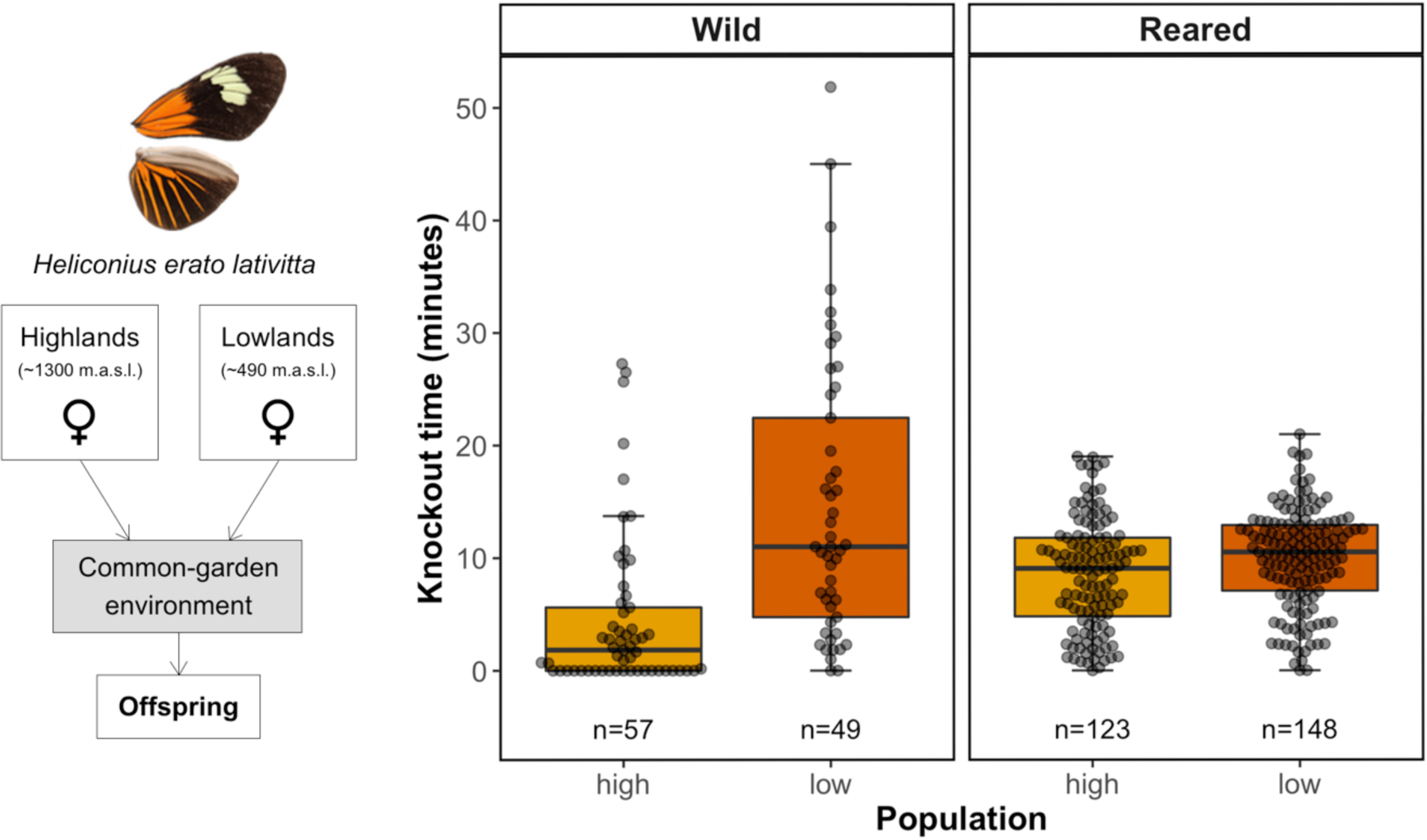
Thermal tolerance in *Heliconius erato*. Experimental design of common garden rearing, fertilised females collected in the highlands and lowlands were brought to a common garden environment and their offspring reared and tested. Heat knockdown time (minutes) across wild individuals (A) and offspring reared in common garden conditions of *Heliconius erato lativitta* populations from high (~489 m.a.s.l., orange) and low elevations (~1344 m.a.s.l., dark orange). Error bars represent 95% confidence intervals of the means. Stars represent significance levels of two sample t-tests between high and low altitude population KO time (*< 0.05, **<0.01, ***<0.001).

## Discussion

We found that the understory of tropical forests had large climatic buffering potential, especially at lower elevations, and that this was similar across two independent elevational clines on both sides of the Ecuadorian Andes (Fig. 1 and Fig. 2). Interpolated climatic variables for these same areas did not capture our observed microclimates, especially at high elevations (~1100 m.a.s.l.), where weather stations are very sparse (Fick and Hijmans, 2017). Furthermore, we found evidence for differences in thermal tolerance in the wild across ten butterfly species, regardless of whether they had altitudinally narrow or widespread distributions (Fig. 4). However, these differences were greatly reduced when a widespread species was reared in common-garden conditions, indicating likely non-genetic, environmental effects on temperature tolerance.

### MICROCLIMATE BUFFERING ACROSS ELEVATIONS

Many macroecological predictions of species distributions have emerged from the notion that tropical latitudes lack strong seasonality (Janzen, 1967; Sheldon et al., 2018; Shelomi, 2012). However, while generally true, these predictions ignore the climate complexity of tropical forests. Our results show that lowland and montane tropical forests buffer ambient temperature and humidity. The understory daily maximum temperature was, on average, 1.4°C and 2.1°C cooler than the subcanopy, at high and low elevations, respectively (Table 1). The temperature offset between forest canopy and understory becomes larger at more extreme temperatures (De Frenne et al., 2019). Thus, lowland forests buffered temperatures to a greater extent than highland forests, which is of particular importance for ectotherms in lowland environments, which are routinely exposed to extreme temperatures (Deutsch et al., 2008). It is important to note that in this study the highland areas were located at mid-elevations and our lowlands were in the foothills of the Andes, which have been less studied because the differences are often assumed to be marginal. Similarly, our subcanopy dataloggers were positioned 10m from the forest floor, thus we would expect the temperature and humidity offsets to be even stronger when compared to above-canopy or open-habitat temperatures.

Janzen’s hypothesis (Janzen, 1967) predicted that the reduced climatic variability in the tropics would result in ectotherms having narrower thermal tolerances, and, in turn, reduce dispersal across elevations (Sheldon et al., 2018). In this study, the mean temperature difference between the subcanopy and the understory, only ten meters apart, was 0.25°C in the highlands and 0.44°C in the lowlands (Table 1). This is more than the temperature change across these elevational clines, where for every 10 m in elevation there was a 0.05°C decrease in temperature (Fig. S9). Yet, in the wild, we found that low altitude populations were much more tolerant to heat than highland populations. While in this study we did not measure cold tolerance, we found that the difference in minimum temperatures across elevations and forest strata is much smaller than that of maximum temperatures (Fig. 1 A5, B5). Thus, we can hypothesize that, given a linear change in cold tolerance, the disproportionally greater heat tolerance of lowland *Heliconius* would result in them having broader thermal breadths than high altitude populations. The microclimatic variability that tropical ectotherms are exposed to within their habitats might offset the lack of seasonality across the year, making some species more able to cope with warming than others. Thus, protecting tropical forests’ climatic buffering potential across elevations is essential to enable potential upland shifting in the face of climate and land-use change.

We found a clear disparity between field-collected microclimate and interpolated macroclimate temperature (Fig. 2). For instance, WorldClim2 estimated the maximum daily temperatures to be 2.01°C higher in the highlands, and overestimated annual mean temperature in the understory by 0.5°C. Strikingly, these values are similar in magnitude to the projected warming for the next century in Andean mid-elevations (Beaumont et al., 2011; Urrutia and Vuille, 2009). In large part, these disparities can be attributed to the fact that coarse-gridded temperature surfaces, such as WorldClim2, are interpolated from weather stations that are located in open habitats (De Frenne et al., 2019; Frenne and Verheyen, 2016). Several recent studies have reported similarly striking differences (Jucker et al., 2018; Lembrechts et al., 2019; Potter et al., 2013; Storlie et al., 2014). Near-surface temperatures can only be accurately measured with in-situ loggers or with emerging remote sensing technologies, such as airborne laser scanning (Jucker et al., 2018). Furthermore, WorldClim2 interpolations at high altitudes tend to be less accurate, especially in the tropics, where weather station data is very sparse (Fick and Hijmans, 2017). This raises the question of how useful coarse macroclimatic grids are for assessing thermal tolerances of organisms which are affected by fine-scale microclimates (De Frenne et al., 2019; Lenoir et al., 2017; Navas et al., 2017; Nowakowski et al., 2018). In addition, very few studies in the tropics have accounted for humidity and vapour pressure deficit variability at the microclimate level (Bujan et al., 2016; Friedman et al., 2019; García-Robledo et al., 2016), as the loggers required to do so can be 4-5 times more costly than temperature loggers. Nevertheless, inclusion of VPD in species distribution models has been shown to significantly improve their accuracy (de la Vega and Schilman, 2017). The differences in VPD between subcanopy and understory observed here were much more pronounced in the lowlands, highlighting the importance of protecting forest complexity in these areas, which are under constant threat of land-use change.

### HEAT TOLERANCE IN THE WILD

Our extremely limited knowledge of thermal tolerance and plastic potential of tropical ectotherms in the wild further hinders our ability to predict ectotherm responses to climate change. As expected, the ability of wild butterfly species to cope with extreme, but natural, levels of heat (~40°C) was much lower for those inhabiting high altitudes. To a lesser extent, highland populations of altitudinally widespread species were similarly intolerant (Fig. 4, Fig. S7). Behavioural shifts might alleviate the impact of climate extremes in tropical forests, but the capacity of a species to shift sufficiently will be constrained by life history, energy budgets, and thermal tolerance (Sheldon and Tewksbury, 2014). In these butterflies, altitude has been shown to pose strong selective pressures, and some are constrained in their body size by contrasting reproductive strategies (gregarious vs. solitary), which could, in turn, restrict adaptive plastic responses to environmental change (Montejo-Kovacevich et al., 2019). Nevertheless, the observed thermal buffering of forests would undoubtedly benefit and be exploited by these butterflies during extreme temperature events, such as the 31 days where temperatures in the lowlands went above 39°C.

### EVIDENCE FOR PLASTICITY IN HEAT TOLERANCE

When reared under a common developmental temperature, the differences in adult thermal tolerance observed in the wild largely disappeared in the widespread species, *H. erato lativitta*, indicating plasticity in individual thermal tolerance (Fig. 4). In this study, we cannot distinguish developmental plasticity from acclimation. The former would imply that larval rearing temperature would irreversibly determine adult heat knock-down resistance, whereas the latter implies a reversible response to temperature (Llewelyn et al., 2018). Interestingly, we did find evidence of at least some genetic component to heat knock-down resistance, as 25% of its variation among common-reared offspring was explained by family identity. Thus, it is likely that a combination of genetic and environmental effects determine adult thermal tolerance in *Heliconius* butterflies. Heat knockdown resistance has been associated with the expression of a heat shock protein (Hsp70) and its plasticity has been shown to vary across altitudinally and latitudinally structured populations of *Drosophila buzzati* and European butterflies (Karl et al., 2009; Luo et al., 2014; Sørensen et al., 2001), suggesting a mechanism for both genetic and environmental control of heat knock-down resistance. Highly heritable thermal performance traits have been shown to allow rapid adaptation to changing climates in lizards (Logan et al., 2014), but developmentally plastic traits with varying and unknown levels of heritability have less predictable roles (Merilä and Hendry, 2014).

The fact that 45% of the individuals from three high altitude *Heliconius* species were knocked out at temperatures between 35-39°C is remarkable (Fig. 3B), given that in the highlands, there were 50 days in a year where temperatures went above 30°C. Canopy-dwelling tropical ants are known to behaviourally circumvent high temperature areas encountered while foraging, avoiding temperatures 5-10°C below their thermal tolerance limits (Logan et al., 2019; Spicer et al., 2017). Thus, these butterflies and other high-altitude species must adapt their flying times and/or behaviours during hot periods, which could have cascading effects on fitness. *Heliconius* follow the same flower foraging trap-lines every day, as they require large amounts of nectar and, uniquely for lepidopterans, pollen, to thrive throughout their long adult life-spans (Jiggins, 2016; Mallet, 1986). Hot patches of the forest close to their thermal limits could severely hinder their ability to follow a foraging path, disrupting a basic biological function. Thus, the climatic refugia that the understory provides could be crucial to cope with the ongoing increase in extreme temperature events (Ruiz et al., 2012; Scheffers et al., 2014).

Relative humidity and vapour pressure deficit (VPD), which have received little attention in the literature, were also greatly buffered by forest canopies (Fig. 3). High VPD has direct impacts on seedling growth and survival (Jucker et al., 2018; Motzer et al., 2005), as well as on ectotherm activity levels (Bujan et al., 2016; Friedman et al., 2019). Nevertheless, forest temperature and humidity buffering may benefit canopy specialists disproportionally. Understory specialist species are thought to be closer to their thermal limits due to the stable climatic conditions they inhabit and thus these may not have suitable refugia or the thermal capacity to cope with warming (Huey et al., 2009; Scheffers et al., 2017; Sheldon, 2019). This could also hold true for less mobile, understory-dwelling life stages, such as butterfly larvae, meaning that these might suffer the greatest detrimental effects on growth and survival under climate change.

### CONSERVATION IMPLICATIONS

Habitat degradation and land-use change in the Andean foothills are pressing concerns for tropical conservation. Ectotherms escaping unsuitable climates in the lowlands may struggle to shift their ranges upwards due to habitat fragmentation (Chen et al., 2009; Scheffers et al., 2016). For example, most remnants of pristine forest in Borneo are in montane areas, as the flat lowlands have been converted to oil palm plantations. Current protected areas may fail to act as stepping stones as they are too isolated and distant from upland climate refugia (Scriven et al., 2015). A similar scenario occurs in Western Ecuador, one of our study areas, where a large portion of the low and mid elevations have been converted to agricultural lands in the past three decades (Wasserstrom and Southgate, 2013). Any habitat change that affects forest heterogeneity could reduce its great temperature buffering potential (Blonder et al., 2018), and butterfly diversity as a whole (Montejo-Kovacevich et al., 2018). Nevertheless, microclimates have been shown to recover decades after low impact land-uses (González del Pliego et al., 2016; Mollinari et al., 2019; Senior et al., 2018), allowing for recolonization of biodiversity (Hethcoat et al., 2019). This highlights the need to protect degraded secondary forest, as these are now more abundant than primary forests worldwide (Edwards et al., 2011; Senior et al., 2017).

## CONCLUSIONS

Tropical ectotherms find themselves in highly heterogenous, threatened habitats, which have greater climate buffering potential than previously thought (De Frenne et al., 2019; Frenne and Verheyen, 2016). However, the low seasonality of tropical environments together with the steep environmental gradients of montane habitats, as Janzen’s hypothesis predicts, makes tropical ectotherms particularly vulnerable to climate and habitat change (Deutsch et al., 2008; Huey et al., 2009; Janzen, 1967; Polato et al., 2018). The clear mismatch between the microclimates we measured and the widely-used interpolated global datasets highlights the importance of field-based climate measurements. Furthermore, the striking difference in heat tolerance and evidence for plasticity across a moderate elevational cline, demonstrates the importance of temperature for the persistence of tropical ectotherms. Our results suggest that the inclusion of microclimate buffering within models and experimental testing of thermal tolerances is crucial for incorporating realistic temperatures experienced by small organisms (Paz and Guarnizo, 2019). More research into the evolvability and plasticity of heat tolerance is needed to accurately assess the vulnerability of tropical ectotherms in the face of anthropogenic change.

## Acknowledgements

We would like to thank all field assistants that have contributed to this study, particularly to Ismael Aldas, Jennifer Smith, and Melanie Brien. We are extremely grateful to the Mashpi Reserve, Narupa Reserve (Jocotoco Foundation), Jatun Satcha reserve, and Universidad Regional Amaóznica Ikiam for their support. Research and collecting permits were granted by the Ministerio del Ambiente, Ecuador, under the *Contrato Marco* MAE-DNB-CM-2017-0058. G.M.K. was supported by a Natural Environment Research Council Doctoral Training Partnership (NE/L002507/1). Funding was provided to C.B. by the Spanish Agency for International Development Cooperation (AECID, grant number 2018SPE0000400194). N.J.N. and C.D.J were supported by the Natural Environment Research Council (grant number: NE/R010331/1).

## Data availability

All data and scripts will be made available in Zenodo upon acceptance (DOI:XXXX, TBC).

